# Long Noncoding RNA Mir17hg and D43Rik Control the Macrophage Response to *Toxoplasma gondii* Infection

**DOI:** 10.1101/2025.01.28.634134

**Authors:** Claire M. Doherty, Jessica Belmares-Ortega, Breanne E. Haskins, Kayla L. Menard

## Abstract

Long noncoding (lnc) RNAs are emerging as imported regulators of immunity but how they impact the host response to infection is still largely unknown. In previous screening studies we identified Mir17hg and D43Rik as host lncRNAs upregulated in mouse bone marrow-derived macrophages regardless of parasite strain type. Here, we examined the impact of Mir17hg and D43Rik during infection of macrophages. Expression of these lncRNAs was transiently increased by *T. gondii*, peaking 6 hr post infection and falling to background noninfected levels by 18 hrs. Both Mir17hg and D43Rik were predominantly localized to the host cell nucleus, suggesting roles in gene regulation. Using a lncRNA silencing approach mediated by siRNA, we uncovered evidence that both D43Rik and Mir17hg were involved in modulating the host transcriptional response to infection with Type I and Type II strains of *Toxoplasma*. These effects were most dramatic during infection with the high virulence Type I RH, where Mir17hg silencing resulted in increased expression of a broad panel of immunity-related genes. Our results implicate lncRNAs Mir17hg and D43Rik as important molecular regulators of the macrophage response to *Toxoplasma*.

## Introduction

The microbial pathogen *Toxoplasma gondii* is a globally distributed intracellular protozoan that infects humans, domesticated animals, and wildlife ^1^. It is estimated to be present in 30% of the global population, and in the United States *Toxoplasma* is a leading cause of foodborne death every year ^2^. In immunocompetent individuals, *T. gondii* establishes a latent infection within the central nervous system where it remains for the lifespan of the host ^3,4^. However, in certain immunocompromised populations, such as HIV-AIDS patients, *T. gondii* may emerge as a pathogen causing life-threatening disease without appropriate therapeutic management ^5^. The parasite also poses a threat during pregnancy, insofar as congenital infection can lead to severe outcomes that include hydrocephaly and abortion of the fetus, as well as sequelae of infection emerging later in life such as blindness, deafness, seizures and mental disability ^6^.

The wide range of outcomes that may result from acquisition of *Toxoplasma* is likely determined in part by interaction of host immunity factors with molecules expressed by the parasite itself ^7^. It is well established that secreted cytokines such as IL-12 and IFN-γ are key to induction of anti-microbial effector cells, while other intracellular host proteins such as members of the IFN-γ-inducible IRG family are important for destruction of the parasitophorous vacuole membrane that harbors intracellular parasites. *Toxoplasma* employs multiple mechanisms to trigger an immune response required for host survival and establishment of parasite latency, while simultaneously adopting other strategies to avoid complete elimination by immune effectors. One such illustration of this principle is the choreographed release of parasite rhoptry and dense granule proteins during host cell invasion and establishment of the parasitophorous vacuole ^8^. For example, dense granule protein GRA24 directly activates p38 MAPK leading to production of IL-12p40 and other proinflammatory cytokines, while rhoptry protein ROP16 triggers STAT3/6 activation leading to an alternatively activated macrophage phenotype and dampening of the T cell response ^9,10^. At the same time, *T. gondii* secretes ROP5 and ROP18 rhoptry proteins that disable IFN-γ-mediated intracellular host killing mechanisms ^11,12^. Maintaining the balance between activation and evasion of immunity likely ensures the success of *Toxoplasma* as a parasitic microbe.

An emerging class of molecules that impact the host-pathogen interaction are non-protein coding (nc)RNAs. It is estimated that only 3% of the human genome is transcribed into proteins, and that up to 80% of the genome is transcribed as ncRNA ^13,14^. One class of non-coding transcripts that are gaining recognition as important immunologically active molecules are long noncoding RNAs (lncRNA), molecules that are defined as possessing a length of ≥ 200 nucleotides ^15,16^. lncRNAs positively and negatively regulate genes at transcriptional and post-transcriptional levels. Functionality, including regulation of gene expression, is largely controlled by RNA secondary structure ^17^. Amongst their possible interactions, lncRNAs can function as gene coactivators, chromatin modifiers, RNA splicing regulators and stabilizers of mRNA ^18,19^. The role of lncRNA in regulation of the immune system is becoming increasingly appreciated ^20^. There is growing interest in exploiting protozoan and helminth lncRNA as therapeutic targets given the ever-growing threat of drug resistance in parasites worldwide ^21^. Indeed, microbial lncRNAs have been identified in the context of both helminth and protozoan parasites ^22–26^. For apicomplexan parasites such as *Plasmodia* spp., *Cryptosporidium* and *Toxoplasma*, a great deal remains to be discovered regarding lncRNAs and their potential to serve as therapeutic targets.

The function of lncRNA in activation and evasion of immunity by *Toxoplasma* is largely unexplored. We previously examined the ability of *Toxoplasma* to trigger host lncRNA responses in infected mouse bone marrow-derived macrophages (BMDM) ^27–29^. RNA-seq analysis of *T. gondii* infected BMDM identified host lncRNA upregulated in a parasite strain-specific manner, as well as other lncRNA responses triggered independently of parasite strain. We hypothesize that host ncRNA responses are another target of manipulation employed by *Toxoplasma*. Recently, we identified two lncRNAs, D43Rik and Mir17hg, that were upregulated during BMDM infection with both Type I (RH) and Type II (PTG) *Toxoplasma* strain types ^27^. While D43Rik is a novel lncRNA, Mir17hg has been identified by an independent group as a lncRNA upregulated in human retinal Müller cells following *T. gondii* invasion ^30^. Here, we focus on the function of these two lncRNA in macrophages. This host cell type was chosen for study because macrophages serve as targets of invasion during in vivo *T. gondii* infection, and because they are important effector cells during the immune response to the parasite ^31^. While bone marrow derived macrophages (BMDM) present many challenges for genetic loss-of-function studies ^32^, we successfully developed an siRNA/ASO RNA silencing approach specifically for this cell type. Using this approach, we found evidence that Mir17hg and D43Rik target a large panel of immunity-related genes.

## Results

### Computational analysis of Mir17hg and D43Rik

lncRNA are generally expressed at very low levels, making downstream analysis difficult ^33^. However, our previous studies identified a relatively high abundance of both D430020J02Rik (D43Rik) and Mir17hg lncRNAs following both Type I- and Type II-*Toxoplasma* infection of BMDM making these ideal targets for further study ^27,28^. The Mir17hg lncRNA is of additional interest because it is conserved in mice and humans ^30^. In mice, D43Rik is located on chromosome 12 and Mir17hg is present on chromosome 14. Both lncRNAs have an exon count of 2, and are similar in nucleotide length (D43Rik: 3,264 nucleotides, Mir17hg: 3,849 nucleotides) (Fig. 1a, b). The Mir17hg lncRNA is expressed from a primary RNA transcript that also encodes a cluster of microRNA species (Mir17-20) that previously were found to be upregulated by *Toxoplasma* ^34,35^ (Fig. 1b). Given the essential role of structure in determining the function of lncRNA ^36^, we generated a predicted optimal secondary centroid structure based on thermodynamic minimum free energy including probable base pairing of each sequence (Fig. 1c and 1d). Total free energy calculated for these structures was -1068.21 kcal/mol for D43Rik and -1186.10 kcal/mol for Mir17hg. These low values indicate high stability and likelihood of the structure prediction ^37^. According to this structure prediction, both lncRNAs have a comparable number of loop structures and possible base pairing interactions along the entirety of the sequence.

**Figure 1.**
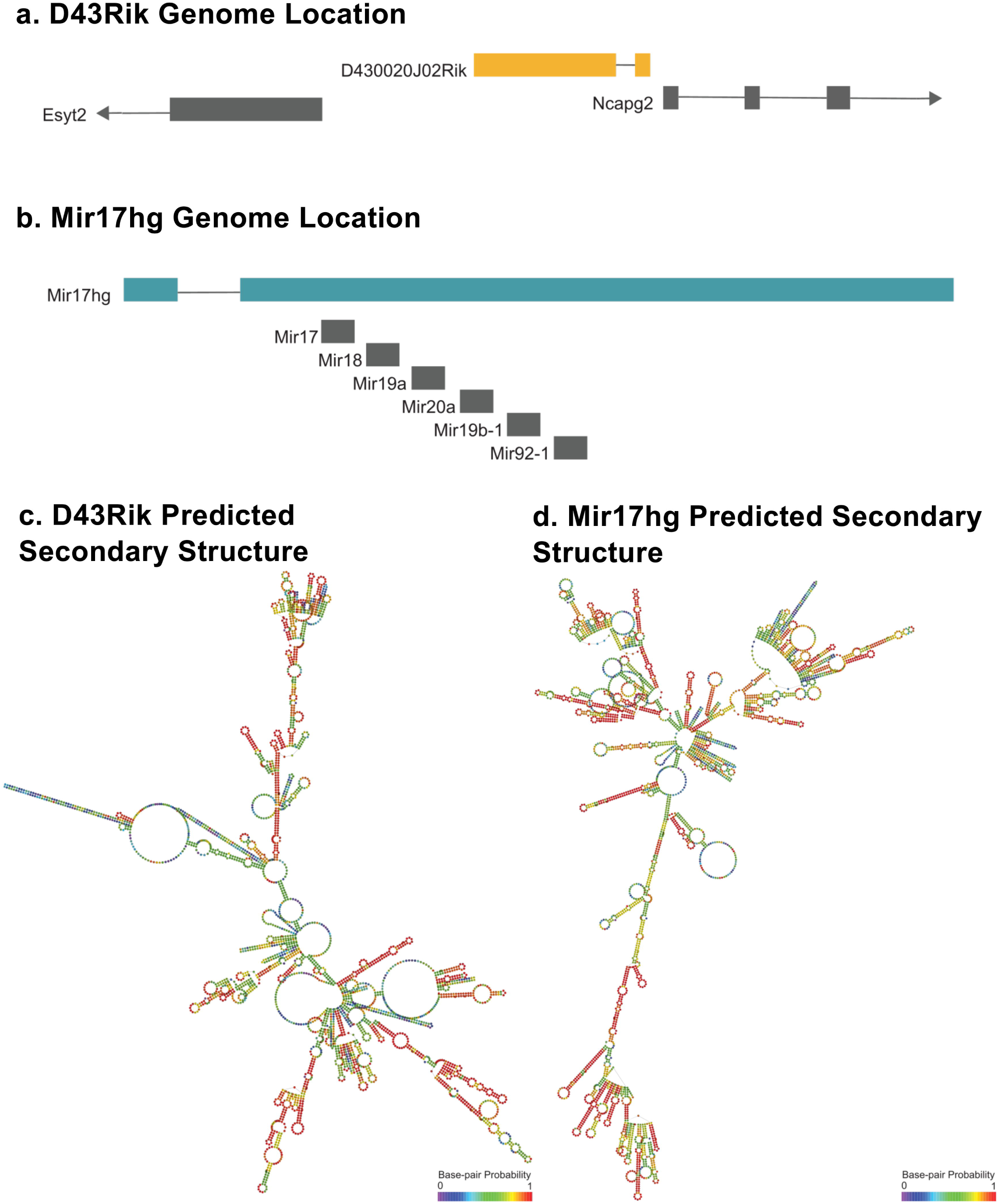
Location and predicted secondary structures of D43Rik and Mir17hg. The genome location of mouse D43Rik (**a**) and Mir17hg (**b**) is shown with neighboring genes and microRNAs respectively. The predicted secondary centroid structures of D43Rik **(c)** and Mir17hg **(d)** were calculated using RNAfold and are based on minimum free energy and base pairing probability.

### lncRNAs Mir17hg and D43Rik localize to the nucleus and are upregulated during *Toxoplasma* infection

Given that lncRNAs may function either within the cytoplasm or the nucleus ^18^, we determined the intracellular location of Mir17hg and D43Rik. Lysates from mouse bone marrow-derived macrophages (BMDM) were separated into cytoplasmic and nuclear fractions then presence of these lncRNAs was examined by RT-PCR analysis (Fig. 2a). In these experiments, we also confirmed the purity of the extracts by examining transcripts encoding GAPDH (cytoplasmic localization) and 7SK (nuclear localization). The results of this study indicate that both D43Rik, and in particular Mir17hg, predominantly localize to the cell nucleus. Next, we determined how expression of Mir17hg and D43Rik changed over time using Type I and Type II *Toxoplasma*. As shown in Fig. 2b, peak levels of both lncRNAs were reached at 6 hr post-infection during Type I infection. Beyond this time point, expression of both Mir17hg and D43Rik rapidly fell to levels present in noninfected BMDM. Type II *Toxoplasma* induced both Mir17hg and D43Rik to similar extent as Type I RH for the first 8 hr (Fig 2d). However, levels of these lncRNA continued to rise in PTG infected cells in contrast to cells infected with RH. A possible explanation for the decrease in lncRNA expression in overnight cultures of RH-infected BMDM was that the cells were simply lysed at this time point. As shown in Fig 2b, this was not the case because BMDM remained intact after 18 hr of infection with both Type I RH and Type II PTG strains. The results instead suggest a parasite strain-specific regulation of these two lncRNA species.

**Figure 2.**
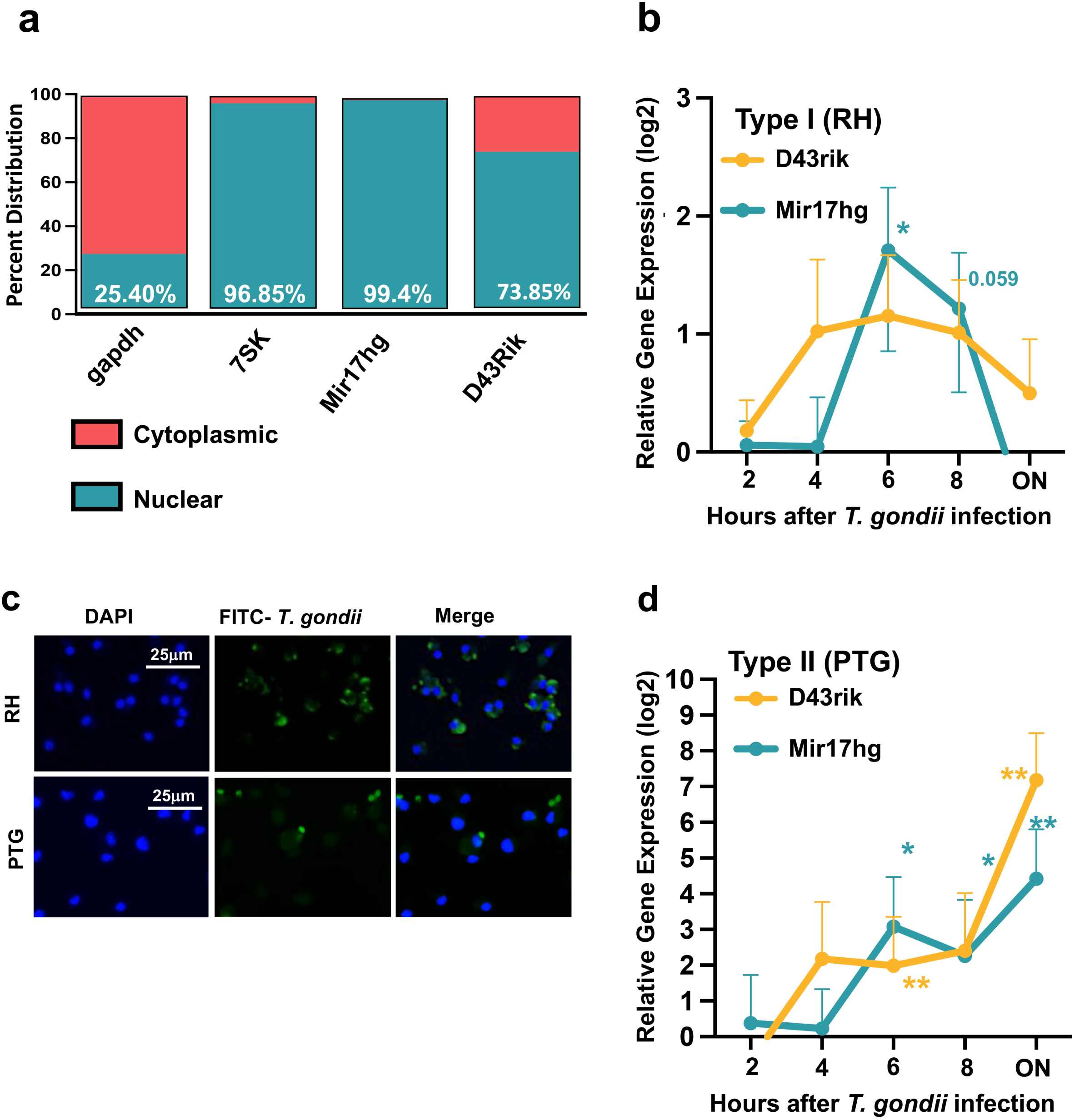
Upregulation of nuclear lncRNA Mir17hg and D43Rik during *Toxoplasma* infection. (**a**) RT-PCR analysis of transcript expression within nuclear and cytoplasmic fractions of bone marrow-derived macrophages (BMDM) reveals localization of MI17hg and D43Rik to the macrophage nucleus. Data represent mean of three independent experiments. (**b**) Kinetics of lncRNA D43Rik and Mir17hg expression during Type I *Toxoplasma* infection and (**d**) Type II *Toxoplasma* infection (MOI 6:1). The data show lncRNA levels relative to expression in noninfected cells. An unpaired Student’s *t* test was used to analyze the data from (b) three or five (d) independent experiments, where **p<0.05 and ***p<0.01. (**c**) Immunofluorescence microscopy of macrophages infected with Type I and Type II *Toxoplasma* for 24 hr and stained with anti-*Toxoplasma* (FITC, green) and DAPI (blue).

### Transcriptional gene silencing in BMDM results in knock down of both lncRNA and protein coding genes

The phagocytic properties of macrophages have posed a challenge in developing an efficient method for delivering oligonucleotides to BMDM through liposome transfection ^38^. Here, we optimized a transfection protocol using Lipofectamine RNAiMAX that results in efficient gene silencing in BMDM of both lncRNA and protein-coding gene transcripts (Fig. 3a). We delivered both siRNAs and anti-sense oligonucleotides (ASO), each of which have unique advantages and disadvantages in gene silencing approaches ^39^. ASOs are particularly useful in lncRNA research, as it they are generally superior to siRNA at targeting nuclear-localized transcripts ^33^. Using ASO targeting different regions of Mir17hg (mir2 and mir4), we achieved a reduction of over 70% of this lncRNA relative to the scramble control in BMDM (Fig. 3b). Similarly, the RNAiMAX protocol using two distinct siRNAs achieved a reduction of over 80% in the relative abundance of lncRNA D43Rik (Fig. 3c). We also tested this protocol for effectiveness in knocking down protein-coding transcripts. Using GAPDH mRNA as a representative example, we silenced expression of this gene by close to 90% (Fig. 3d).

**Figure 3.**
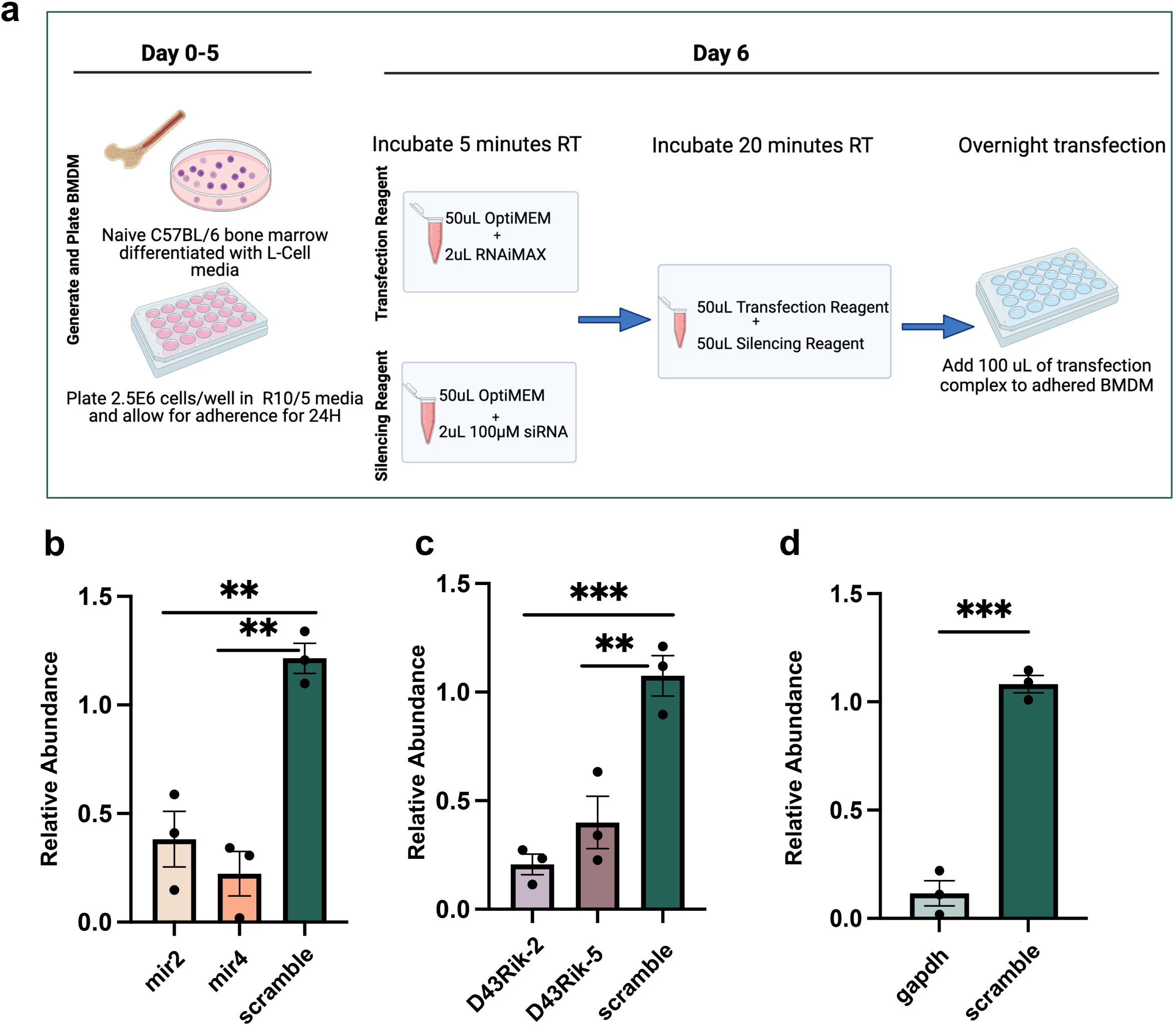
siRNA transfection of BMDM functionally silences Mir17hg and D43Rik lncRNA and protein-coding GAPDH mRNA expression. (**a**) Schematic diagram of the siRNA transfection design. BMDM were generated according to established protocols in L cell media. On Day 5, cells were plated into 96 well plates in R10/5 media and allowed to adhere for an additional 24 hr. The silencing reagents (anti-sense oligonucleotides for mir17hg and siRNA for D43Rik) were added to OptiMEM and incubated for 5 minutes at room temperature, followed by combination with Lipofectamine RNAiMAX and incubation for 20 min at room temperature. The resulting complexes were then added to adhered BMDM for 18-24 hours of transfection. (**b**) Relative abundance of mir17hg in BMDM following transfection with anti-sense oligonucleotides (ASO) mir2 and mir4 that target mir17hg along with ASO scramble control. (**c**) Relative abundance of D43Rik following transfection with siRNAs D43Rik-2 and D43Rik-5 along with siRNA scramble control. (**d**) Relative abundance of mRNA following transfection with GAPDH-targeting siRNA. The graphs in panels **b-d** show relative abundance compared to non-transfected cells. An unpaired Student’s *t* test with Welch’s correction was used to analyze the data from three independent experiments, where **p<0.05 and ***p<0.01.

### Mir17hg acts as a regulator of host immunity during *Toxoplasma* infection

With the ability to silence these lncRNAs in macrophages, we turned our focus to understanding their contributions to the immune response during *Toxoplasma* infection. Using a commercial PCR array that probes for 84 immunity-related genes, we first assessed the impact of RH and PTG *T. gondii* infection in the absence of lncRNA silencing. Type I *Toxoplasma* (Fig. 4a) infection induced a larger panel of inflammation-related transcripts than observed for Type II *Toxoplasma* (Fig. 4b). Parasite strain-specific effector molecules are well-established modulators of host immunity ^40^, and such effector molecules are likely to underlie the differences we found here.

**Figure 4.**
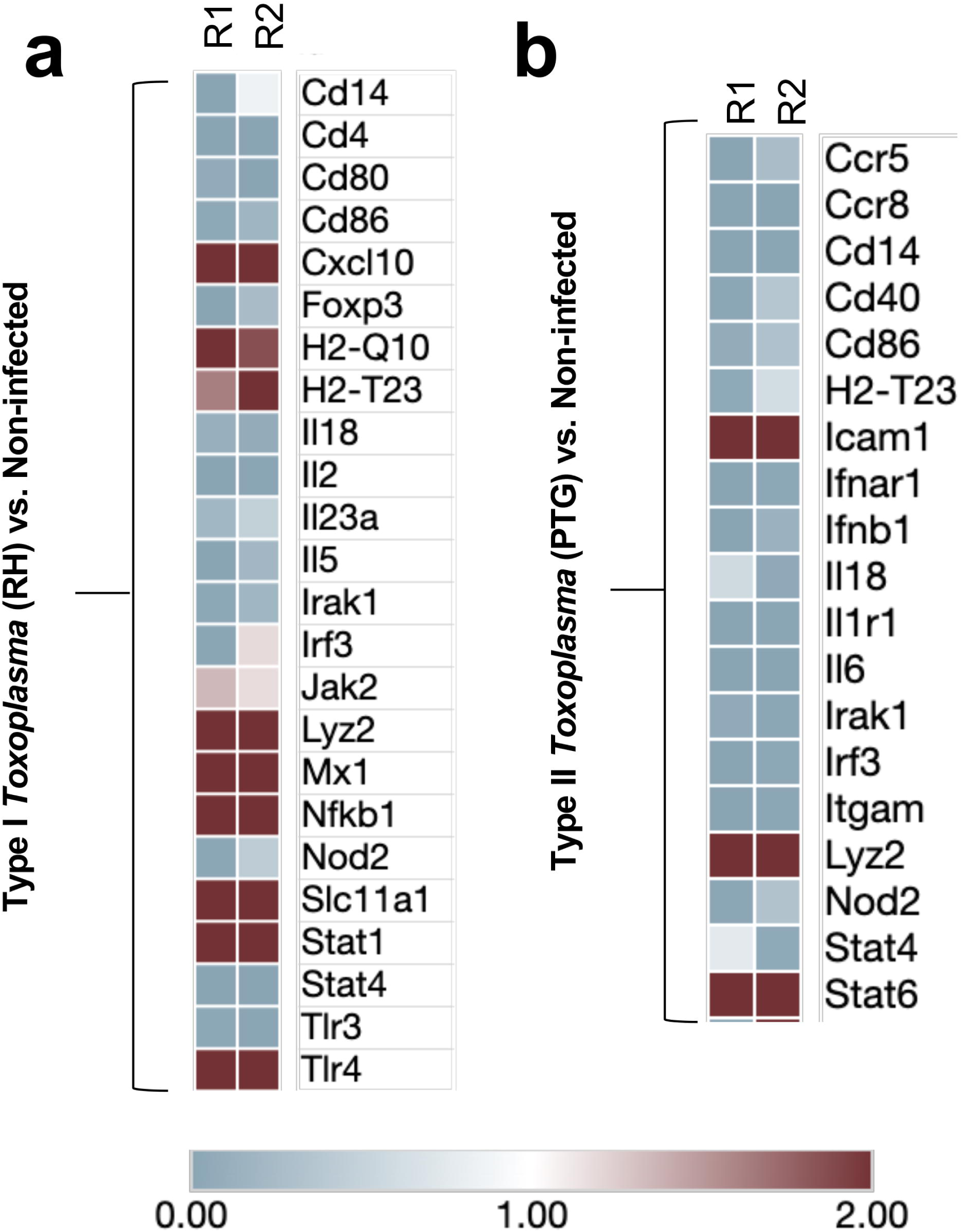
*Toxoplasma* infection modulates macrophage immunity-related genes in strain-specific manner. (**a**) RT-PCR differential gene expression of immunity-related genes in macrophages infected with Type I *Toxoplasma* (RH, MOI 6:1) relative to noninfected controls. (**b**) Differential gene expression of infected BMDM (Type II-PTG, MOI 6:1) relative to noninfected controls. The data in the two panels show two independent experiments (R1, R2). Heatmaps were generated using *Morpheus* software provided by the Broad Institute with relative gene expression standardized around min=0 and max=2. In these studies, parasite-positive cells constituted 70-90% of the population as determined by immunofluorescence microscopy.

RelA (p65) is a key subunit of the NFκB transcription factor that regulates many immunity genes. While very little is known about Mir17hg, one recent study implicated a role for this lncRNA in upregulation of *relA* in human colorectal cancer cells ^41^. Accordingly, we sought to determine how this lncRNA affected *relA* in noninfected and infected BMDM. Silencing of Mir17hg using siRNA led to a significant increase in *relA* abundance in BMDM suggesting that Mir17hg acts as a negative regulator of RelA (Fig. 5a). We also determined the effect of Mir17hg knockdown in cells harboring parasites. The enhancing effect of Mir17hg knockdown was maintained in cells infected with *T. gondii* (Fig. 5a). The contrasting result of the effect of this lncRNA in our study and that of other investigators may reflect differences in cell types and host species in each case. We also note that *Toxoplasma*-induced Mir17hg upregulation and consequent *relA* downregulation does not inherently contradict previous data showing Type II GRA15 upregulation of NFκB ^42^. This is because the latter work focuses on activation of RelA whereas the present study address expression levels of this transcription factor.

**Figure 5.**
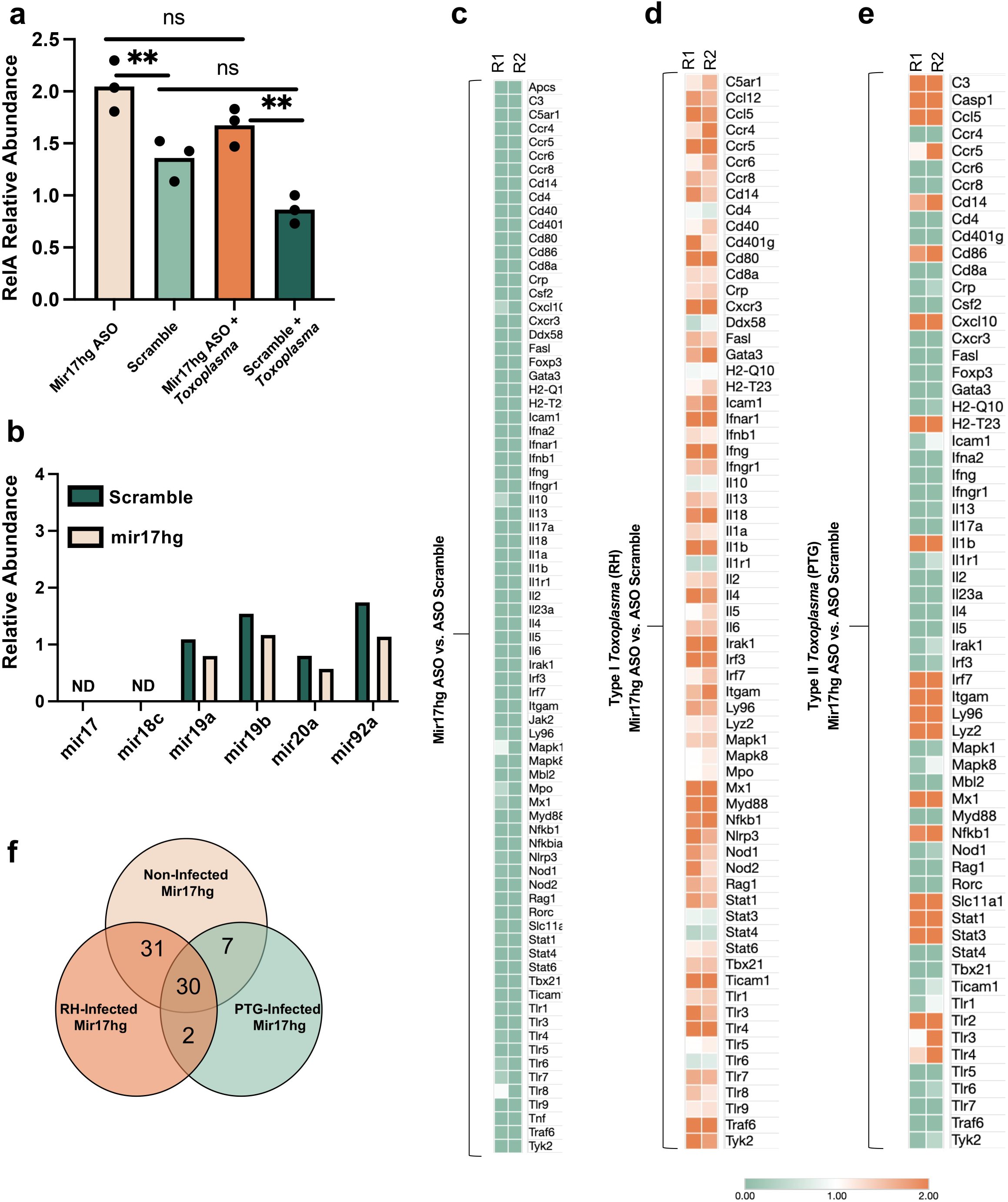
Silencing of Mir17hg results in widespread effects on immune gene expression. (**a**) Relative abundance of *relA* gene expression following knockdown of mir17hg along with respective ASO scramble control, mir17hg + *T. gondii* infection, and ASO scramble + *T. gondii* infection. Data are expressed relative to nontransfected cells. Three independent multiple unpaired *t* test were used to analyze the data where **p<0.01. (**b**) Relative abundance of microRNAs neighboring lncRNA mir17hg following mir4/scramble siRNA transfection compared with nontransfected controls. One representative experiment of three independent replicates; nd, non-detected. Data are expressed relative to nontransfected cells. (**c**) RT-PCR differential gene expression of immunity-related genes in noninfected BMDM with mir17hg knocked down relative to scramble control. (**d**) Differential gene expression of infected BMDM (Type I-RH, MOI 6:1) under mir17hg knockdown conditions relative to infected ASO scramble BMDM. (**e**) Differential gene expression of infected BMDM (Type II-PTG, MOI 6:1) under mir17hg knockdown conditions. The data in c-e show two independent experiments (R1, R2). Heatmaps were generated using *Morpheus* software provided by the Broad Institute with relative gene expression standardized around min=0 and max=2. In experiments employing infected BMDM, parasite-positive cells constituted 70-90% of the population as determined by immunofluorescence microscopy. (**f**) Number of differentially expressed immune genes shared between non-infected BMDM, RH-infected BMDM, and PTG-infected BMDM after Mir17hg knockdown, representing the data visualized in panels (c-e).

The lncRNA Mir17hg is expressed as a primary RNA transcript with several microRNAs (mir17-20) that possess immunoregulatory properties (Fig. 1b) ^43^. To ensure that the transfection protocol targeting lncRNA Mir17hg did not inadvertently affect these microRNAs, the relative abundance of the mir17-92 cluster was assessed under Mir17hg knockdown relative to scramble conditions. The results showed no detectable difference in microRNA expression after siRNA transfection compared to scramble control for mir19a, mir19b, mir20 and mir92a (Fig. 5b). We also note that that the mir17-92 cluster was expressed at consistently low levels throughout our study. The lack of significant difference between the scramble control and transfected samples suggests that this microRNA cluster is not impacted by ASO targeting of lncRNA Mir17hg.

To further investigate the role of lncRNA Mir17hg in immune regulation, a commercial PCR array was employed to assess the impact of Mir17hg silencing on 84 immune-related genes. The findings revealed a general dampening of immune-related genes relative to the scramble control following Mir17hg knockdown (Fig. 5c). We speculate that this might reflect an impact of Mir17hg on the final stages of BMDM when ASO are added. Notably, when Mir17hg was knocked down in the context of infection with the highly virulent RH strain of *Toxoplasma*, there was an overall upregulation of key immune genes compared to the infected BMDM treated with the scramble control (Fig. 5d). In particular, we found upregulation of several genes for chemokines/chemokine receptors (*ccl12*, *ccl5*, *ccr4*, *ccr5*, *cxcr3*) as well as IL-1 family cytokines (*Il1b*, *Il18*). In the case of infection with the less aggressive Type II PTG strain, silencing of Mir17hg had more diverse effects on host immunity genes (Fig. 5e). For example, while several transcripts of interest were strongly upregulated (*casp1*, *ccl5*, *irf7*, *stat1*, *stat3*), roughly equal numbers were downregulated (including *ccr6*, *ccr8*, *myd88*, *traf6*). The Venn diagram (Fig. 5f) summarizes the results shown in Fig. 5c-e. We identified 30 immune genes that were affected by the silencing of Mir17hg expression in all treatments, suggesting the significance of this lncRNA in modulating key immune genes both in noninfected and infected macrophages (Fig. 5f). Remarkably, only two immune genes exhibited differential expression exclusively in parasite infected BMDM. Together, these results suggest an important role for lncRNA Mir17hg in modulating macrophage immune responses, particularly in the context of *Toxoplasma* infection.

### Knockdown of D43Rik has diverse effects on key immune genes during *Toxoplasma gondii* infection

D43Rik is flanked by two genes, *esty2* and *ncapg2* (Fig. 1a). To determine whether siRNA knockdown of D43Rik also affected the expression of these neighboring genes, their relative abundance was evaluated in D43Rik siRNA samples and siRNA scramble controls (Fig. 6a). There was no significant difference in the expression of the neighboring genes, indicating that the silencing protocol was specific D43Rik lncRNA itself. We then examined how D43Rik silencing affected immunity genes in noninfected and infected BMDM. For unclear reasons, the results were notably more variable than those obtained with Mir17hg knockdown (Fig. 6b-d). However, we note that during RH infection several gene transcripts were upregulated by D43Rik silencing (examples include *ccr4*, *ccr8*, *cd40*). In BMDM infected with PTG, many more genes were downregulated in D43Rik silenced cells (for example, *ccl12*, *ccl5*, *Il23a*, *nod1*, *nod2*). These collective findings emphasize the parasite strain-dependent nature of immune responses during infection and suggest that immune manipulation mediated by D43Rik and Mir17hg has both positive and negative effects on expression of immunity related genes. Although the influence of D43Rik is less pronounced than that of Mir17hg, we identified 13 genes that exhibited differential expression upon D43Rik silencing, regardless of the treatment conditions (Fig. 6e). Moreover, while the overall contribution of D43Rik in immune gene expression is less dramatic in comparison to Mir17hg, there was a more evident association with differential gene expression during *T. gondii* infection, with 14 genes showing differential expression exclusively in the infected samples (Fig. 6e).

**Figure 6.**
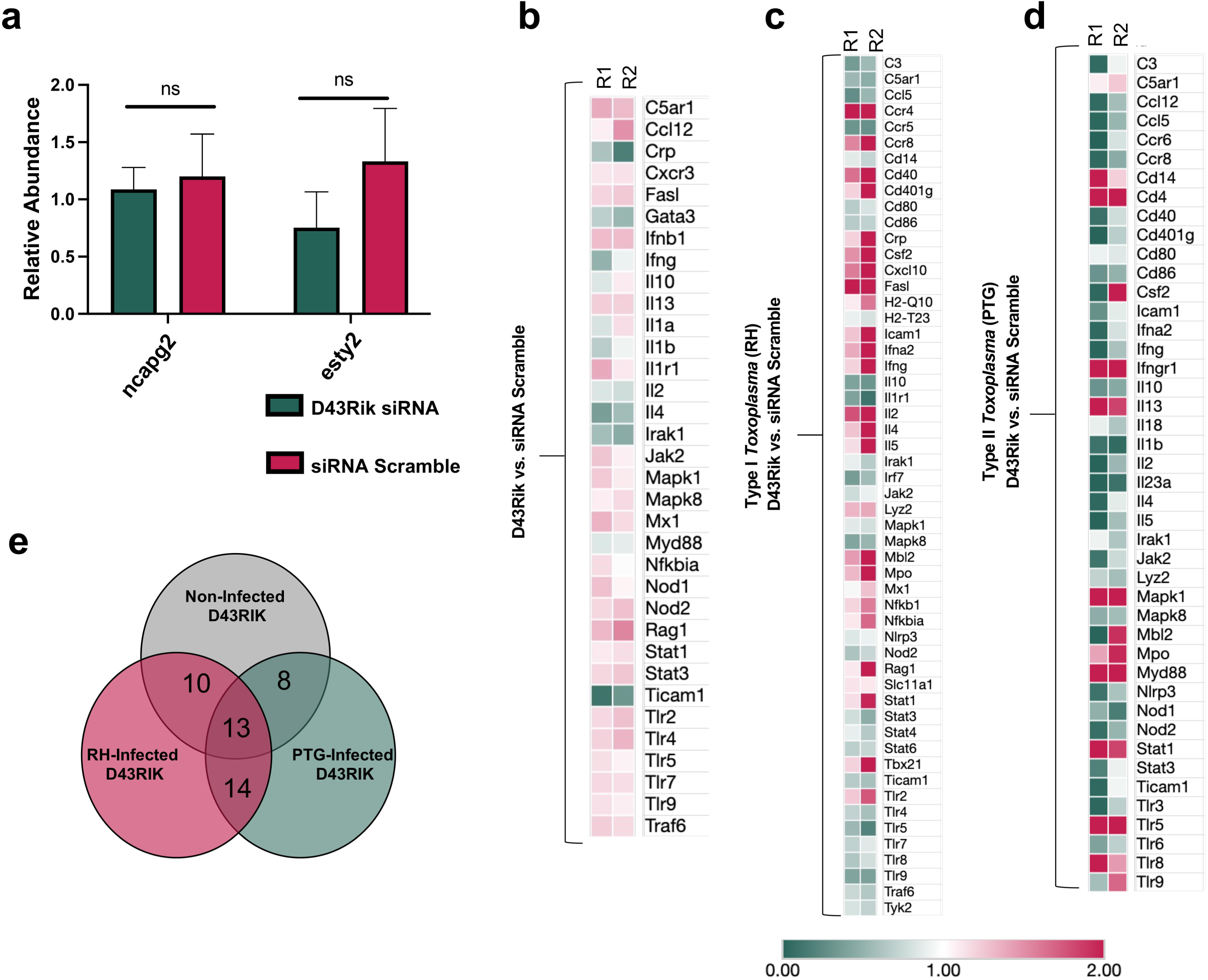
siRNA knock down of D43Rik results in effects on immune gene expression without impacting expression of flanking genes. (**a**) Relative abundance of genes neighboring D43Rik following silencing of D43Rik along with siRNA scramble control compared to nontransfected cells. Three independent experiments were performed and multiple unpaired *t* tests were used to analyze the data. ns = non-significant. (**b**) RT-PCR differential expression of immune related genes in BMDM with D43Rik knocked down relative to scramble control. (**c**) Differential gene expression of infected (Type I-RH, MOI 6:1) BMDM under D43Rik silencing conditions relative to infected BMDM with no siRNA knockdown. (**d**) Differential gene expression of infected BMDM (Type II-PTG, MOI 6:1) under D43Rik knockdown conditions relative to infected siRNA scramble. In panels b-d, heatmaps were generated using *Morpheus* software (Broad Institute) with relative gene expression standardized around min=0 and max=2. Two independent experiments are displayed. In these experiments, parasite-positive cells constituted 70-90% of the population. (**e**) Number of differentially expressed immune genes shared between non-infected BMDM, RH-infected BMDM, and PTG-infected BMDM after D43Rik knockdown, representing the data visualized in panels (b-d).

### Influence of Mir17hg and D43rik on *T. gondii* infection in BMDM

Given the impact of Mir17hg on immunity-related genes during *T. gondii* infection, we further examined the influence of lncRNA silencing on the infection itself. Upon knockdown of Mir17hg in BMDM, we observed a significant increase in infection at 24 hr post-inoculation. The increased parasite infection was evident in both Type I (Fig. 7a and Fig. 7b) and Type II (Fig. 7c and Fig. 7d) infections. This was somewhat unexpected given the overall upregulated inflammatory profile observed when Mir17hg was silenced (Fig. 5d, e). Nevertheless, it is possible Mir17hg knockdown has either direct or indirect effects on the host cell metabolism that results in a more favorable environment for the parasite ^44^. In contrast, silencing D43Rik expression had no observable effect on infection with either Type I (Fig. 7e and Fig. 7f) or Type II (Fig. 7g and Fig. 7h) *T. gondii*. These findings highlight the critical regulatory role of Mir17hg in modulating immunity-related macrophage genes during *T*. *gondii* infection and suggest an impact of this lncRNA on parasite infection.

**Figure 7.**
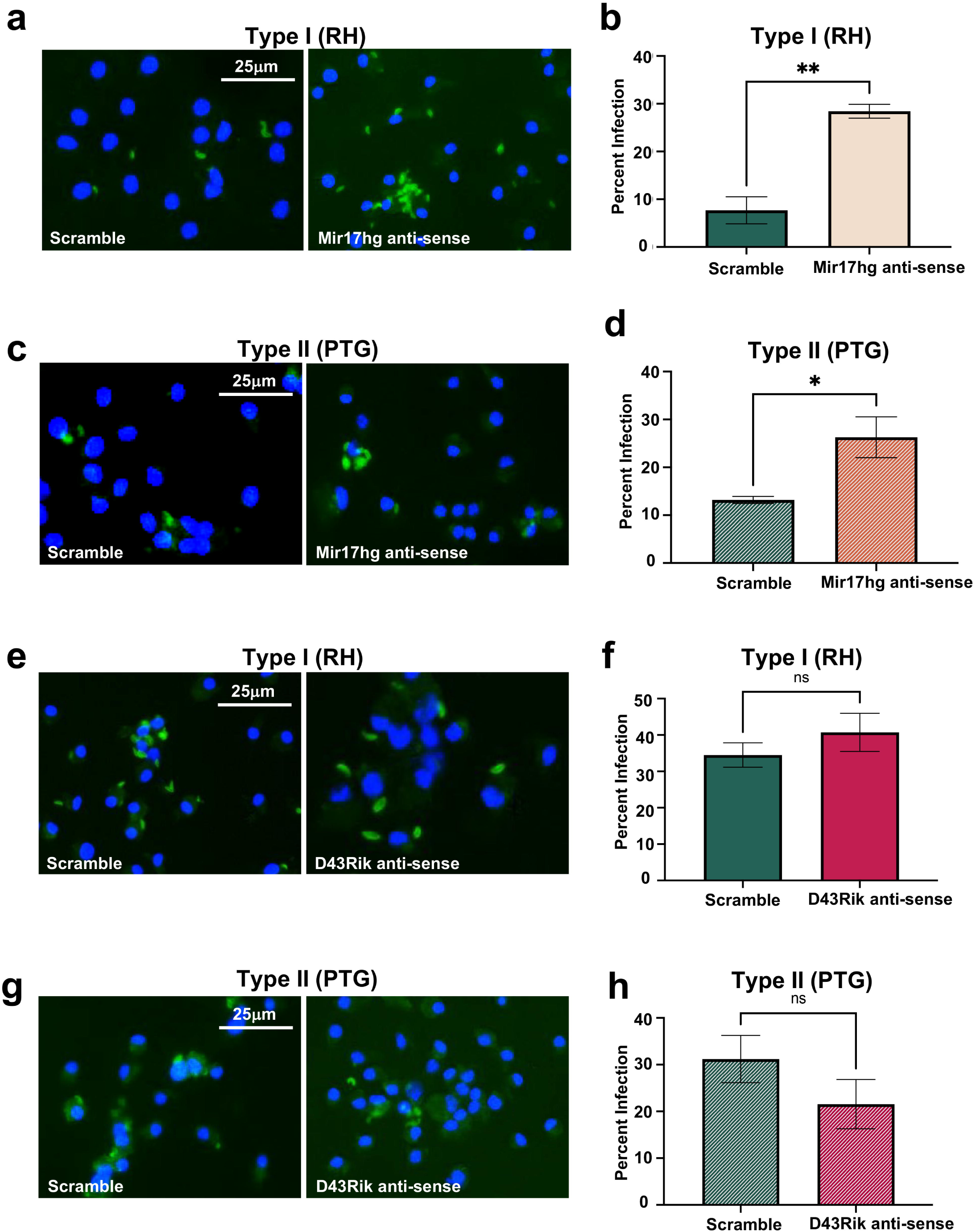
Effect of Mir17hg and D43Rik silencing on *Toxoplasma* infection in macrophages. (**a**) BMDM infected with Type I *T. gondii* (RH) under Mir17hg transfection conditions and ASO scramble control at 24 hours post-infection; cells were permeabilized and stained with antibody to *T. gondii* (FITC, green) and nuclei were stained with DAPI (blue). (**b)** Quantification of Type I *Toxoplasma* infected BMDM with silenced Mir17hg and scramble controls. (**c**) BMDM infected with Type II *T. gondii* (PTG) under Mir17hg transfection conditions and ASO scramble control at 24 hours post-infection along with corresponding quantification (**d**). (**e**) BMDM infected with Type I *T. gondii* (RH) with D43Rik silenced via siRNA transfection along with siRNA scramble controls along with corresponding quantification (**f**). (**g**) BMDM infected with Type II *T. gondii* (PTG) under D43Rik silencing alongside quantification (**h**). All infections were carried out at an MOI of 6:1. Data are representative of three independent experiments. An unpaired Student’s *t* test was used to analyze three fields of view with a total of 100-200 cells. **p<0.05 and ***p<0.01.

## Discussion

Macrophages and macrophage-related cells play a key role in the host-parasite interaction during encounter with *Toxoplasma*, serving as preferential targets of infection, producers of key cytokines, and acting as microbicidal effector cells ^31^. Here, we used BMDM to examine the role of host lncRNA during *T. gondii* infection. We selected for study D43Rik and Mir17hg, lncRNAs previously reported to be upregulated during Type I and Type II *Toxoplasma* infection, for further analysis ^27,30,45^. Both D43Rik and Mir17hg strongly localized to the host cell nucleus, suggesting functions associated with transcriptional rather than post-transcriptional regulation. Using an siRNA knockdown approach, we uncovered evidence that D43Rik and Mir17hg lncRNAs target a large panel of immune-related transcripts. For example, siRNA knockdown of Mir17hg in BMDM resulted in dampening of immune-related genes relative to the scramble control RNA. In the context of infection, however, the silencing of Mir17hg resulted in upregulation of a large panel of key inflammatory immune genes encoding chemokines and chemokine receptors (for example, *ccl12*, *ccl5*, *ccr4*, *ccr5*, *cxcr3*) as well as IL-1 family cytokines (*Il1b*, *Il18*). Despite similar infection levels, this striking upregulation of inflammatory genes was predominantly observed in the virulent Type I RH strain. Nevertheless, knock down of Mir17hg during infection with the less virulent Type II PTG strain resulted in increased expression of inflammatory genes (*casp1*, *ccl5*, *irf7*, *stat1*, *stat3*) coincident with decreased expression of other inflammatory genes (*ccr6*, *ccr8*, *myd88*, *traf6*). Thus, despite the fact that both parasite strains upregulate Mir17hg, the impact of this lncRNA on host gene expression differed in the context of *Toxoplasma* strain (Type I vs. Type II). A possible explanation for this finding is that Mir17hg lncRNA interacts with additional ncRNAs that themselves are expressed in a parasite strain-specific manner. Previous RNA-seq work from our group identified a subset of 33 lncRNAs that are expressed in a strain-specific manner during Type I RH and Type II PTG infection, providing a list of candidate noncoding RNAs to examine ^27^. Similar parasite strain-specific trends were observed in infected BMDM in which D43Rik expression was knocked down, although for unclear reasons responses were notably more variable than those following Mir17hg silencing.

We observed up-regulation of NFκB family member *relA* after Mir17hg silencing, suggesting this lncRNA acts as a transcriptional repressor of NFκB expression. The RelA (p65) protein forms a dimer with p50 that activates both proinflammatory and anti-apoptotic genes upon release from its endogenous inhibitor, IκB ^46^. Other lncRNAs are known to mediate both positive and negative regulation of macrophage NFκB involving transcriptional and post-transcriptional mechanisms ^47^. For example, HOX transcript anti-sense intergenic RNA (HOTAIR) is required for degradation of IκB during NFκB activation ^48^. Another lncRNA, metastasis-associated lung adenocarcinoma transcript 1 (MALAT1), binds to p50/p65 preventing its transactivating function ^49^. Interestingly, in models of hepatocellular carcinoma, lncRNA 00607 acts as a tumor suppressor by binding to the *rela* promoter thereby blocking gene activation ^50^. In this regard, lnc00607 resembles Mir17hg insofar as both function as negative regulators of *rela* expression. We note another group reported that Mir17hg upregulates *relA* transcription by acting as a sponge for an inhibitory micro-RNA in the context of colorectal cancer ^41^. The disparate result between that study and ours might be a consequence of the different cell types used (transformed colorectal cancer cells vs. non-transformed BMDM). Regardless, it is interesting to note that in our experiments, many of the genes upregulated upon Mir17hg silencing in infected cells are dependent upon NFκB signaling.

Silencing of Mir17hg resulted in a notably broad upregulation of immunity-related genes that was particularly striking during RH infection. It is tempting to speculate a chromatin-modifying repressive function for Mir17hg insofar as other lncRNA molecules are known to have important effects on chromosomal epigenetic remodeling ^51,52^. For example, HOTAIR binds to approximately 800 sites in the human genome where it imparts repressive histone signatures through interaction with polycomb repressive complex-2 (PRC2) ^53^. Even more profoundly, the lncRNA Xist initiates epigenetic suppressive changes that culminate in silencing of the entire female X chromosome ^54^. It seems possible that upregulation of Mir17hg expression by *Toxoplasma* is a broadly acting evasion mechanism employed by the parasite to mute intracellular responses to infection.

There were striking differences in macrophage gene expression during infection with Type I RH relative to Type II PTG under lncRNA knockdown conditions. Because infection rates were comparable, these effects are likely due to the secretion of polymorphic parasite host-directed effector molecules from rhoptry or dense granule organelles ^55^. Two prime candidates are ROP16 and GRA15. The rhoptry kinase ROP16_I/III_ activates STAT3/5/6 leading to an overall M2 macrophage gene expression profile ^56–58^. In contrast, dense granule protein GRA15_II_ interacts with TNF receptor-associated factors resulting in NFκB activation and a general M1 macrophage phenotype ^57,59^. Several lncRNA have been identified as regulators of M1/M2 differentiation that could be involved in ROP16/GRA15 mediated macrophage polarization ^47^. We are currently investigating the role of these parasite effector molecules under conditions of Mir17hg and D43Rik silencing.

The observation of a significant increase in parasite infection along with increased levels of inflammatory transcripts under Mir17hg knockdown conditions requires further study. The results presented here show that Mir17hg functions as a regulator of host infection regardless of parasite strain type and associated strain-specific effectors such as ROP16 and GRA15. How Mir17hg impacts the outcome of in vivo *Toxoplasma* infection awaits the creation of appropriate lncRNA gene knockout mouse models.

Initially regarded simply as transcriptional noise in the host cell, it is now understood that lncRNAs are an integral component in regulation of cellular activity. However, their well-recognized cell specificity as well as general lack of conservation across species make lncRNAs a challenge to study ^15^. Recently, CRISPR methods for BMDM have been developed using CRISPR-Cas-9 and sgRNA ^60^. While effective for protein-coding genes, sgRNA induce frameshift or nonsense mutations, which are generally not effective for loss-of-function studies in lncRNA ^33^. Other CRISPR methods have been successfully utilized for lncRNA (such as two paired gRNAs or targeting splice sites). However, these would need to be adapted for BMDM and may be problematic for lncRNAs that overlap protein-coding genes or miRNAs (as is the case with mir17hg). Overall, there is likely value in employing both CRISPR and traditional RNA interference methods for studying lncRNA in BMDM. In conclusion, how host lncRNA responses contribute to immunoprotection or immunopathology during infection, and how microbial pathogens such as *Toxoplasma* exploit these responses to promote persistence are areas ripe for investigation.

## Materials and Methods

### Ethics

All experiments were performed in strict accordance with the recommendations set forth by the National Institutes of Health *Guide for the Care and Use of Laboratory Animals (8^th^ Edition).* Protocols were approved by the Institutional Animal Care and Use Committee at the University of New Mexico (Animal Welfare Assurance Number A4023-01). All efforts were made to minimize animal suffering and distress over the course of the studies.

### Mice

C57BL/6 mice were obtained from The Jackson Laboratory. Both male and female mice (6-12 weeks of age) were used in these studies. Animals were housed and cared for under an IACUC approved protocol (22-201240-MC). The mice were euthanized using CO_2_ asphyxiation, as recommended by the American Veterinary Medical Association. All authors complied with ARRIVE guidelines,

### Parasites

High virulence Type I strain (RH) and low virulence Type II strain (PTG) *Toxoplasma* were purchased from the American Type Culture Collection (ATCC) and were maintained *in vitro* by passage on confluent monolayers of human foreskin fibroblasts (ATCC). Parasite cultures were tested for *Mycoplasma* contamination every 6 months (MycoProbe Detection Assay; R & D Systems, Minneapolis, MN).

### Cell Culture

Bone marrow derived macrophages (BMDM) were generated from tibias isolated from C57BL/6 mice. Bone marrow single cell suspensions were collected using a 10-mL syringe and a 27-Ga needle and differentiated into bone marrow-derived macrophages using media composed of Dulbecco’s modified Eagle media (BMDM; Cat#10-017-CV), 10% bovine growth serum (BGS; HyClone Laboratories, Cat# SH30541.03), nonessential amino acids (ThermoFisher Scientific, Cat#11140-050), 0.1 mg/mL streptomycin (ThermoFisher Scientific, Cat #15140-122), and L929 culture supernatant as previously described ^61^. The fully differentiated BMDM were collected for studies on day 5 of culture.

### Immunofluorescence Microscopy

Differentiated BMDM were plated in BMEM containing 10% BGS, 100 U/mL penicillin, and 0.1 mg/mL streptomycin on glass coverslips overnight (37C, 5% Co2). Following a 5:1 infection with RH or PTG, coverslips were collected and processed for microscopy at the timepoints indicated. After rinsing with PBS, cells on coverslips were fixed in 3.7% paraformaldehyde (MilliporeSigma, Cat #FX0410-5) for 20 min at room temperature. Following fixation, coverslips were blocked for 1h at room temperature using 5% normal mouse serum (Invitrogen, Cat #10410) in permeabilization buffer (0.1% saponin in PBS). Staining was accomplished in permeabilization buffer using fluorescein isothiocyanate (FITC)-conjugated anti-*Toxoplasma* polyclonal antibody (Invitrogen, Cat #PA17253). Coverslips were mounted onto microscope slides using Permount solution containing 4’,6-diamidino-2-phenylindole (DAPI; Invitrogen, Cat #P36962). Imaging was performed using a BX53 fluorescence microscope (Olympus America, Inc) and DP manager software (Olympus). Each slide was imaged over 5 different fields each containing approximately 100 DAPI-positive cells.

### Quantitative RT-PCR

Total RNA was prepared using a RNeasy Mini Kit (Qiagen, Cat #74004), and the samples were subjected to Turbo DNase treatment (Invitrogen, Cat #AM2238). RNA was converted to cDNA using the SuperScript IV VILO Master Mix (Invitrogen, Cat# 11756050). Quantitative PCR was performed on target genes and normalized to the expression of the housekeeping gene *Ppia* using the SYBR green method (SsoAdvanced Universal SYBR Green Supermix, Bio-Rad, Cat #1725275) and the Bio-Rad CFX96 RT-PCR machine. Expression relative to uninfected control samples was calculated using the ΔΔCt method. A control with no added reverse transcriptase was included for each sample. In some cases, uninfected samples were occasionally at the limit of detection of the qRT-PCR assay. In these instances, 37 cycles as the Cq (quantification cycle). Primer sequences employed for amplification are shown in Supplementary Fig. 1.

### Nuclear and Cytoplasmic Fractionation

Nuclear and cytoplasmic fractions were prepared from BMDM using the NE-PER Nuclear and Cytoplasmic Extraction Reagents (ThermoFisher Scientific, Cat #78835). RNA was isolated, reverse transcribed and target transcripts amplified as described above for quantitative PCR. In these experiments presence of GAPDH and 7SK transcripts served as controls for purity of cytoplasmic and nuclear extracts, respectively. The percentage of target in each fraction was calculated by dividing the population of each fraction by the sum of the nuclear and cytoplasmic populations.

### Computational Analysis

Mouse lncRNA sequences were downloaded and visualized by UCSC Genome Browser (GENCODE VM32: D43Rik ENSMUST00000220477.2, Mir17hg ENSMUST00000134140.3). A thermodynamic ensemble prediction of each lncRNA was obtained using the RNAfold webserver based on parameters as described ^62^. Briefly, the minimum free energy (MFE) and partition function fold algorithm were selected to calculate an optimal secondary structure. A predicted centroid secondary structure and base pairing probability of the output was visualized through the same program.

### Transfection of BMDM

Day 5-differentiated BMDM were plated at a concentration of 2.5 x 10^5^/well in 24 well tissue culture plates in 400 uL of RPMI 1640 with supplemental glutamine, 10% FBS, 5% L-cell culture media (R10/5 media). BMDM were allowed to adhere to plates overnight (5% CO2, 37°C). Transfection was accomplished by combining Lipofectamine RNAiMax (Invitrogen Cat# 13778-030), silencing reagent, and OptiMEM (Invitrogen Cat# 31985-070). For every reaction, 50 µL OptiMEM and 2 µL RNAiMax were combined and incubated for 5 minutes RT. Concurrently, 50 µL OptiMEM was combined with 2 µL of the appropriate silencing reagent at a 100 µM concentration and allowed to incubate for 5 minutes RT. Following this brief incubation, both components were combined in equal parts and allowed to incubate for 20 minutes at RT with care taken to not physically disrupt the reagent. 100 µL of the resulting mix was added to the adherent macrophages in 400 µL of R10/5 media. Transfections commenced 18-24 hours prior to downstream studies. Antisense oligonucleotides were used to knock down mir17hg (Antisense LNA GapmeR Standard, Qiagen, Cat# 339511) along with ASO negative scramble control (Qiagen, Cat# 339515). Small interfering RNA (siRNA) reagents were used to silence D43Rik (Silencer Select siRNA, Invitrogen, Cat# 4390771) along with corresponding siRNA negative control scramble (Silencer Select Negative Control No.1 siRNA, Cat# 4390843).

### Qiagen Arrays

RT^2^ Profiler PCR arrays (Mouse Innate & Adaptive Immune Responses; Qiagen Cat# PAMM-0527D-12/330231) were accomplished following manufacturer’s instruction. RNA collection and cDNA synthesis was performed as described in detail above.

### MicroRNA abundance

MicroRNA was collected using the miRNeasy Tissue/Cells Advanced Mini Kit (catalog #217604) according to manufacturer’s instructions. cDNA synthesis targeting microRNA was accomplished using the miRCURY LNA RT Kit (catalog #339340). miRCURY LNA SYBR Green PCR Kit (Qiagen, Cat# 339346) was used for qPCR detection of the amplification products. qPCR allowed for the assessment of the abundance of relevant microRNA. Primers for microRNA are as follows: hsa-mir17-5p hsa-mir18a-5p, hsa-mir19a-3p, hsa-mir19b-3p, has-mir20a-5p, mmu-mir92a-3p, 5S rRNA (catalog #339306).

## Acknowledgement

Supported by grants from the US National Institute of Allergy and Infectious Diseases (EYD, AI139628; EYD, AI170657).

## Data availability statement

All data generated or analyzed during this study are included in this published article.

## Author contributions

CMD and KLM designed and performed the experiments, analyzed data and proof-read the manuscript. BEH and JBO performed experiments, analyzed data and proof-read the manuscript. EYD and CMD drafted the manuscript. EYD designed the experiments and analyzed the data.

