## Supplementary figures and images for "Long Noncoding RNA Mir17hg and D43Rik Control the Macrophage Response to *Toxoplasma gondii* Infection"

### Supplemental Table 1

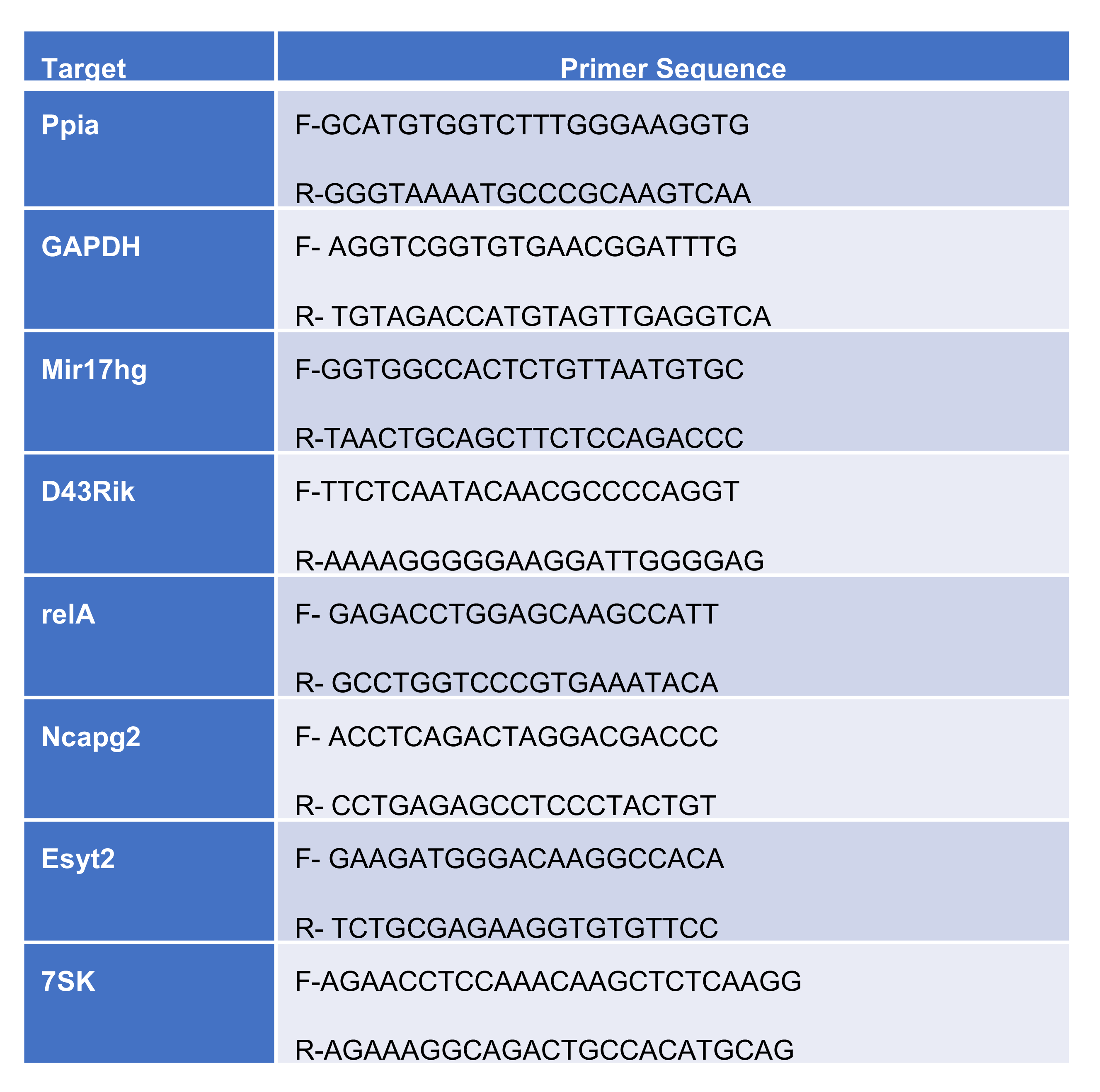
